# Exploration of bacterial bottlenecks and *Streptococcus pneumoniae* pathogenesis by CRISPRi-seq

**DOI:** 10.1101/2020.04.22.055319

**Authors:** Xue Liu, Jacqueline M. Kimmey, Vincent de Bakker, Victor Nizet, Jan-Willem Veening

## Abstract

*Streptococcus pneumoniae* is a commensal bacterium of the human nasopharynx, but can cause harmful infections if it spreads to other parts of the body, such as pneumonia, sepsis or meningitis. To facilitate pathogenesis studies, we constructed a doxycycline-inducible pooled CRISPR interference (CRISPRi) library targeting all operons in protypical *S. pneumoniae* strain D39V. Our library design allows fitness within the pool to be assessed by a one-step PCR reaction directly followed by Illumina sequencing (CRISPRi-seq). The doxycycline-inducible CRISPRi system is tightly controllable and suitable for both bottleneck exploration and evaluation of gene fitness *in vitro* and *in vivo*. Here, we applied CRISPRi-seq to identify genetic factors important for causing pneumococcal pneumonia. Mice were infected intratracheally with our CRISPRi library and bacteria collected at 24 h (from lung) and 48 h (from both lung and blood) post-infection. CRISPRi-seq showed a critical bottleneck at 48 h after intratracheal infection, with only a few bacteria surviving the brunt of the innate immune response to cause systemic infection. However, earlier at 24 h post-infection, many significant differences in gene fitness cost between *in vitro* and *in vivo* conditions were identified, including genes encoding known and putative novel virulence factors, genes essential only *in vivo*, and genes essential only *in vitro*. A key advantage of CRISPRi-seq over traditional transposon-based genetic screens is that all genes, including essential genes, can be tested for their role in virulence and pathogenicity. The approaches developed here should be generally applicable to study infection bottlenecks and *in vivo* fitness for other important human and animal pathogens.

## Introduction

*Streptococcus pneumoniae* is one of the most prevalent opportunistic human pathogens. The bacterium frequently colonizes the human nasopharynx but can cause severe diseases when it invades normally sterile sites. Invasive pneumococcal diseases, including pneumonia, sepsis, and meningitis, lead to millions of deaths per year (Weiser et al., 2018). *S. pneumoniae* is the leading agent of bacterial pneumonia worldwide (van der Poll and Opal, 2009). The pathogenesis of pneumococcal pneumonia involves complicated host-pathogen interactions, and while several key virulence factors are well studied, it remains unknown if or how the majority of the bacterium’s genome contributes to disease progression. Murine *S. pneumoniae* infection is commonly studied for modeling clinically relevant stages of disease including pneumonia and sepsis (Chiavolini et al., 2008). High-throughput identification of important pneumococcal factors during the progression of bacterial pneumonia in the murine model can provide new perspectives for understanding this leading human infectious disease.

Large-scale identification of *S. pneumoniae* virulence determinants has been attempted by signature-tagged mutagenesis (STM) and Tn-seq studies (Chen et al., 2007; Hava and Camilli, 2002; Lau et al., 2001; Opijnen and Camilli, 2012; van Opijnen et al., 2009), however, these approaches have certain technical limitations, including the inability to investigate essential genes, and the fact that not all Tn-insertions result in gene inactivation thus requiring large libraries to fully cover the genome. We previously harnessed an IPTG-inducible CRISPRi system for functional study of essential genes in *S. pneumoniae* D39V *in vitro*, and with an arrayed CRISPRi library, we refined the essential gene list and identified the function of several hypothetical proteins (Liu et al., 2017). This prior study showed the power of CRISPR interference (CRISPRi) for functional gene analysis; however, it was laborious to handle the arrayed library and the IPTG-inducible system was not ideal for *in vivo* studies, limiting its application scope. Based on these considerations, we developed a tetracycline/doxycycline-inducible CRISPRi system for *S. pneumoniae* that is applicable to both *in vitro* and *in vivo* studies. In addition, we constructed a pooled CRISPRi library targeting nearly all operons of the prototypic *S. pneumoniae* strain D39V (Slager et al., 2018) that can readily be combined with Illumina sequencing (herein referred to as CRISPRi-seq). This sgRNA library also covers core operons of other pneumococcal strains, like R6 (estimated 87.9% of genetic elements covered), TIGR4 (75%), Hungary 19A-6 (72.3%), Taiwan 19F-14 (72.2%), 11A (69.6%), G54 (72.5%) (see supplementary text). While pooled CRISPRi libraries have recently been reported for *Escherichia coli, Staphylococcus aureus, Vibrio natriegens* and *Mycobacterium tuberculosis* (Cui et al., 2018; Jiang et al., 2020; Lee et al., 2019; Wang et al., 2018; de Wet et al., 2018), they all used large pools of sgRNA targeting each gene multiple times, often leading to off targeting (Cui et al., 2018). In addition, these libraries required very deep sequencing to obtain enough statistical power on the abundance of each sgRNA in the population and are thus not well suited for conditions in which bottlenecks appear. Here, we carefully selected a single sgRNA for every operon in *S. pneumoniae* D39V, thereby limiting off target effects and reducing the pool of sgRNA required to cover the entire genome.

In the present study, we used a murine pneumonia model initiated by intratracheal infection with our pooled library, followed by CRISPRi-seq to measure the relative fitness contribution of each operon in the *S. pneumoniae* D39V genome. In addition to identifying genes associated with bacterial survival in the lung and blood niches, we also identified an extreme bacterial bottleneck in progression from pneumonia to sepsis. Bottlenecks limit population diversity, reminiscent of earlier work in which a small effective population of pneumococcal nasopharyngeal colonization was observed (Li et al., 2013). In addition, prior reports provided a hint regarding a single-cell bottleneck for bloodstream invasion and transmission, however these studies lacked resolution as they utilized only three isogenic mutants (Gerlini et al., 2014; Kono et al., 2016). In contrast, our pooled CRISPRi library contains 1499 different genetic markers coded by the various sgRNAs allowing precise measurement of bottlenecks. Our studies further demonstrate a large variation between hosts, and ultimately in disease progression – even in genetically identical inbred mice. Specifically, while an extreme bottleneck to bloodstream infection was observed in each mouse, the number of individual unique bacterial “barcodes” detected in blood ranged from 0 – 10^7^ in each mouse. This finding is significant to our understanding of pneumococcal disease, given that in humans, the majority of *S. pneumoniae* exposures *do not* lead to severe disease, and disease manifestations can vary within a host over time. Thus, these high-resolution studies allow tracking of bacterial dynamics within a host, a first step towards demonstrating that no gene is singularly responsible for “virulence” or “avirulence.” Rather, it appears likely that there are many possible combinations through which the individual genes of the pneumococcal genome confer pathogenicity.

## Results

### A tetracycline-inducible CRISPRi system in *S. pneumoniae* enables both *in vitro* and *in vivo* studies

To enable the study of *S. pneumoniae* genes *in vivo*, we designed a tetracycline inducible (tet-inducible) CRISPRi system. The two key elements of the CRISPRi system, *dcas9* and *sgRNA* were integrated into the pneumococcal chromosome, driven by a tet-inducible promoter (P_tet_) and a constitutive promoter (P3), respectively (Figure 1A). Constitutively expressed *tetR*, which encodes the tet-repressor, was codon optimized and integrated into the chromosome to enable tet-inducible expression conferred by P_tet_ (Sorg et al., 2019). Here, doxycycline was used to induce dCas9 expression because it has been extensively validated as tet inducer in rodent models due to its high potency and excellent tissue penetration (Redelsperger et al., 2016). To alleviate growth stress caused by doxycycline’s antimicrobial activity, TetM served as the antibiotic marker for *dcas9* chromosome integration. TetM is a ribosome protection protein that confers tetracycline resistance by catalyzing the release of tetracycline and its derivatives from the ribosome in a GTP-dependent manner (Burdett, 1996; Dönhöfer et al., 2012).

**Figure 1.**
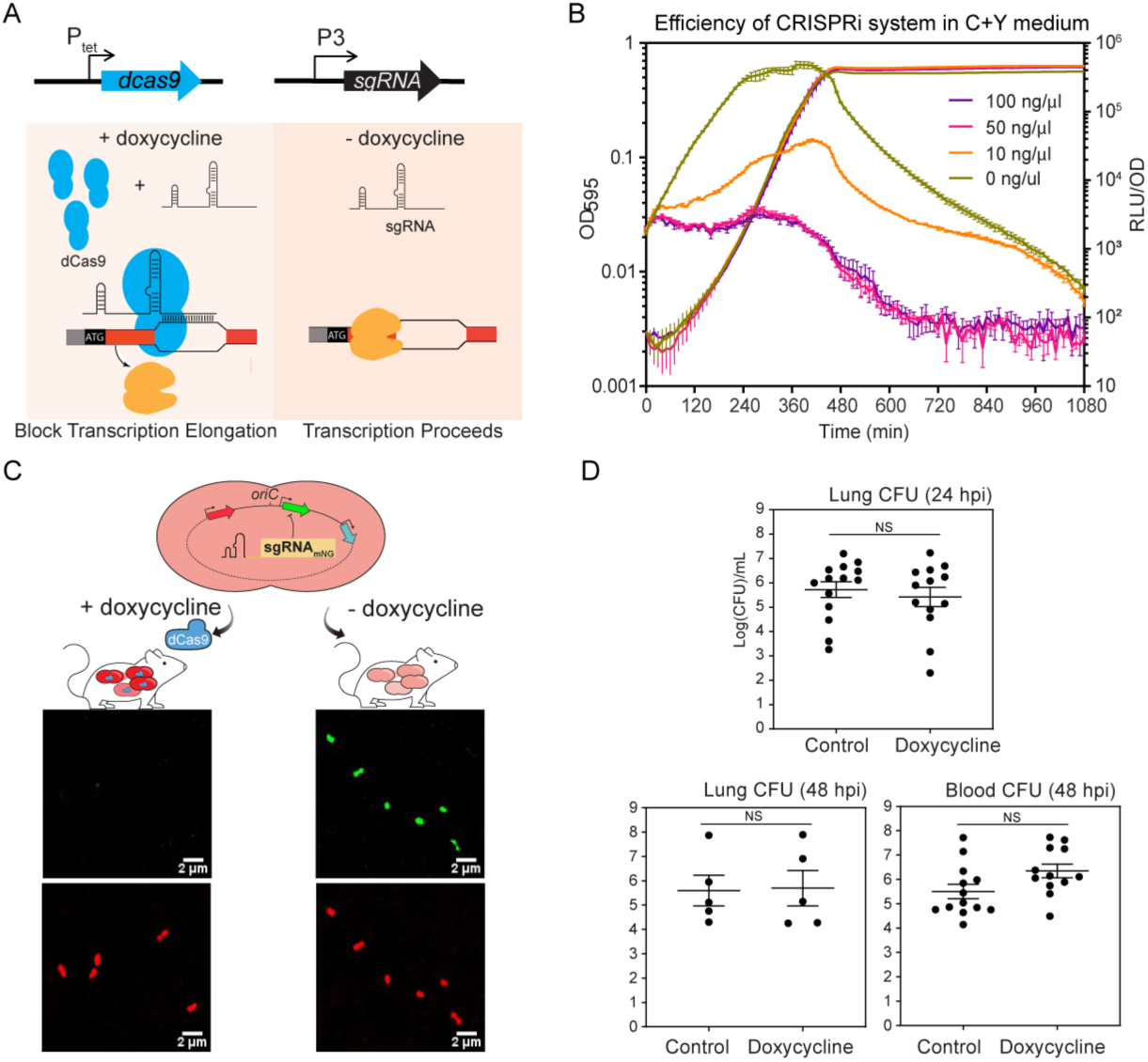
A tet-inducible CRISPRi system in *S. pneumoniae*. (A) The design of the tet-inducible CRISPRi system. The two key elements, *dcas9* and *sgRNA*, were integrated into the chromosome and driven by a tet-inducible promoter (P_tet_) and a constitutive promoter (P3), respectively. With addition of the P_tet_ inducer, here doxycycline, a derivative of tetracycline, dCas9 is expressed and binds to the target under the guidance of a constitutively expressed sgRNA. The specific dCas9-sgRNA binding to the target gene acts as a transcriptional roadblock. P_tet_ is tightly controlled in the absence of the inducer, and expression of dCas9 is repressed. Without binding of dCas9-sgRNA, the target gene is transcribed. (B) The CRISPRi system was tested by targeting a constitutively expressed reporter gene, *luc*, which encodes firefly luciferase. The system was induced with doxycycline at different concentrations. Luciferase activity (RLU/OD) and cell density (OD595) were measured every 10 minutes. The values represent averages of three replicates with SEM. (C) Reporter strain to assess *in vivo* activity of the doxycycline-inducible CRISPRi system. Strain VL2351 constitutively expresses two fluorescent proteins, mNeonGreen and mScarlet-I, in which mNeonGreen is targeted by the sgRNA. Bacteria were collected from blood of mice on control or doxy-chow at 48 hpi and imaged with confocal microscopy in both the red and green channels. (D) Bacterial load at both lung and blood was enumerated by plating on agar plate. Each dot represents a single mouse. Mean with SEM was plotted. There is no significant (NS) difference between the bacterial load in control and doxycycline treated mice (Mann Whitney test).

The efficiency of the tet-inducible CRISPRi system was tested in both C+Y medium and in a mouse pneumonia model. A reporter strain expressing firefly luciferase (*luc*) under a constitutive promoter was used for the *in vitro* assay performed in C+Y medium. Efficiency of the tet-inducible CRISPRi system targeting *luc* was evaluated by monitoring luciferase activity in C+Y medium with different concentrations of doxycycline (Figure 1B). As little as 10 ng/ml doxycycline was enough to strongly reduce (>20 fold) luciferase activity within 3 h, while 50 ng/ml doxycycline yielded a maximum repression efficiency without causing growth retardation (Figure 1B). To test the functionality of the tet-inducible CRISPRi system *in vivo*, a dual-fluorescent reporter strain was constructed that constitutively expressed mNeonGreen and mScarlet-I. The CRISPRi system in this reporter strain was designed with an sgRNA targeting the coding region of mNeonGreen. BALB/c mice were fed *ad libitum* with chow containing 200 ppm doxycycline hyclate or control chow for 2 days prior to infection, then infected with the reporter strain by intratracheal inoculation. At 48 h post infection (hpi), bacteria in blood samples were checked by confocal microscopy for both green (mNeonGreen) and red (mScarlet-I) fluorescence. As expected, both mNeonGreen and mScarlet-I signals were present in the sample harvested from mice fed with control chow, whereas the mNeonGreen signal was absent in mice receiving doxycycline (Figure 1C). Specific inhibition of *S. pneumoniae* mNeonGreen expression in mice fed with doxycycline confirmed functionality of the tet-inducible CRISPRi system *in vivo*. Finally, we verified that this dose of doxycycline did not alter *S. pneumoniae* burden in blood or lungs at 24 hpi and 48 hpi (Figure 1D), giving us a tool to regulate gene expression without interfering with bacterial survival.

### A concise CRISPRi library with 1499 sgRNAs targeting the entire genome of *S. pneumoniae* D39V

Due to the well-documented polar effects inherent to a CRISPRi system (Bikard et al., 2013; Liu et al., 2017; Peters et al., 2016; Qi et al., 2013) we adopted this technique to study gene function at the operon level. The transcriptional units of *S. pneumoniae* D39V are well annotated in our previous study (Slager et al., 2018) (https://veeninglab/com/pneumobrowse). First, for each operon led by a primary transcriptional start site (pTSS), i.e. the only or strongest TSS upstream of a feature (Slager et al. 2018), one specific sgRNA was selected with preference for a short distance downstream from the TSS (Figure 2A), resulting in 794 unique sgRNAs. However, the pTSS operons cover only about 65% of the genetic elements of *S. pneumoniae* D39V. For the genes not covered within a pTSS operon, one specific sgRNA was selected to target the coding region closest to the start codon, leading to the design of 705 further sgRNAs (Figure 2A). In total, 1499 sgRNAs were selected to target 2111 out of 2146 genetic elements of *S. pneumoniae* D39V. 35 elements are not included due to lack of a protospacer adjacent motif (PAM) or localization in repeat regions on the chromosome (supplementary table S1). Potential (off-)targets of sgRNAs are listed in supplementary table S2.

**Figure 2.**
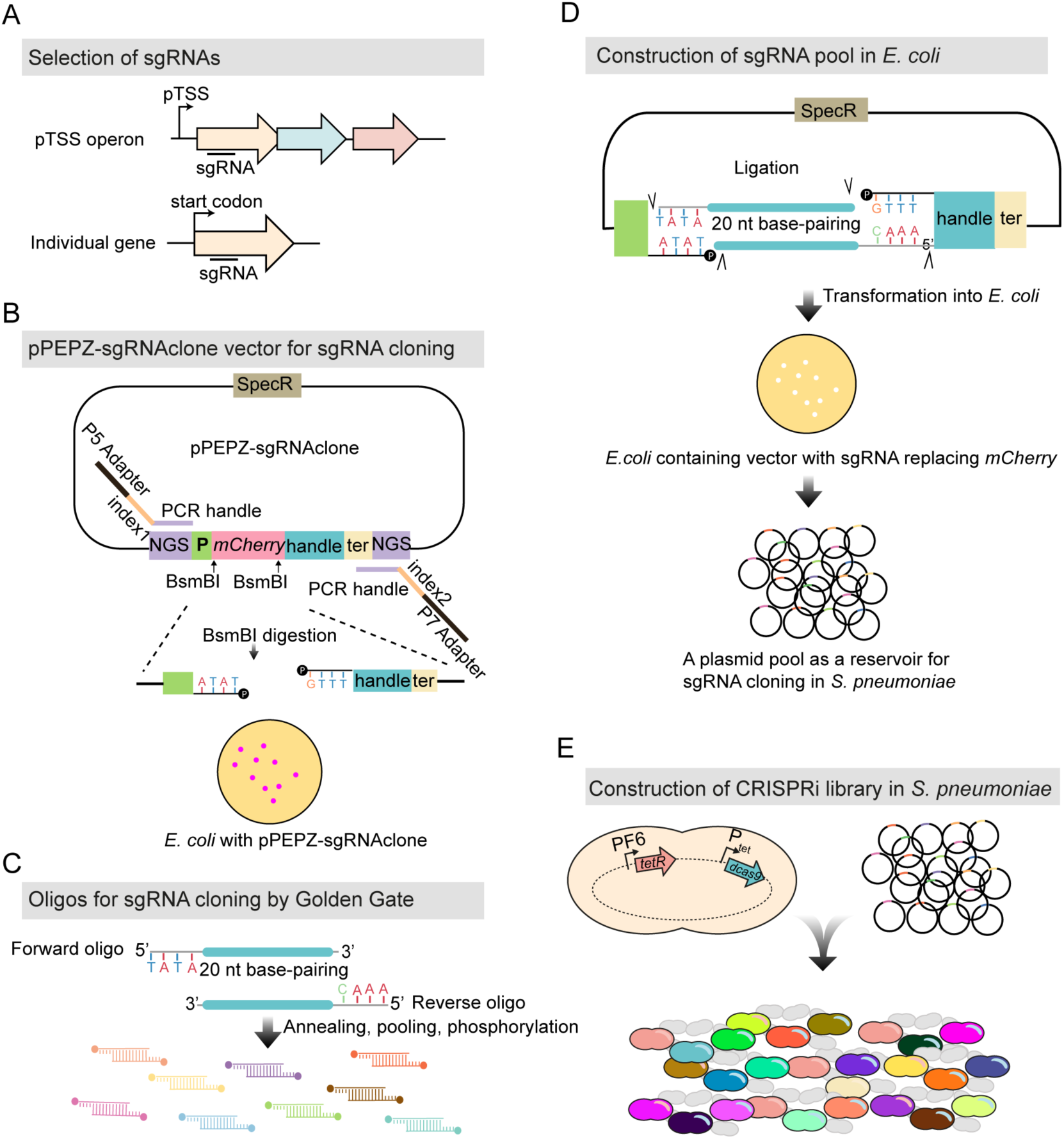
Workflow for construction of the pooled tet-inducible CRISPRi library. (A) For the operons driven by primary transcriptional start site (pTSS) (Slager et al., 2018), an sgRNA in proximity of the pTSS was selected, resulting in 794 sgRNAs. For the genes not covered by pTSS operons, an sgRNA in proximity of the start codon was selected, resulting in 705 sgRNAs. In total, 1499 sgRNAs were selected targeting 2111 genetic elements out of the 2,146 in *S. pneumoniae* D39V. (B) The vector for sgRNA cloning, named pPEPZ-sgRNAclone, was designed to enable high efficiency Golden Gate cloning, monitoring false positive ratio, and construction of Illumina library in a one-step PCR. SpecR is the spectinomycin resistant marker; NGS indicates key elements which allow construction of Illumina library by one-step PCR (see methods); P is the constitutive promoter which drives the expression of sgRNA; *mCherry* encodes a red fluorescent protein placed in the base-pairing region of sgRNA and flanked by a BsmBI site on each end; handle and ter represent the dCas9 handle binding region and terminator of sgRNA. *E. coli* with the pPEPZ-sgRNAclone form red colonies resulting from the expression of mCherry. BsmBI digestion of the vector produces ends that are compatible with the sgRNA oligo annealing in (C). (C) Forward and reverse oligos were designed for each sgRNA containing 20 bp complementary to sgRNA and 4 nt overhangs compatible with the BsmBI digested vector. The oligos were annealed and pooled together followed by 5’ phosphorylation. (D) Ligation product of the digested vector (B) with the sgRNA annealing (C) was transformed into *E. coli*. *E. coli* transformed with the vector containing the sgRNA show white colonies due to replacement of *mCherry* with *sgRNA*. Among the more than 70,000 colonies (provides about 50 fold theoretical coverage of the sgRNA diversity) we collected, no red colony was observed. The collected colonies were pooled together, and plasmids purified from the *E. coli* pool serves as an sgRNA reservoir. (E) Pooled plasmid library was transformed into a *S. pneumoniae* D39V strain with chromosomally integrated tet-inducible *dcas9*.

To enable efficient and convenient construction of sgRNA libraries, we engineered a new vector, pPEPZ-sgRNAclone, that allows sgRNA insertion via Golden Gate cloning (Figure 2B). The *mCherry* reporter, flanked by a BsmBI site on each end, was inserted at the position of the base-pairing region of the to-be-cloned sgRNA. *E. coli* containing this parent vector produce red colonies on agar plates due to mCherry expression (Figure 2B). To facilitate Illumina sequencing, key Illumina elements, including read 1, read 2, and adaptor sequences, were inserted flanking the sgRNA transcriptional unit. A forward oligo and reverse oligo were synthesized encoding each designed sgRNA. Each oligo was 24-nt, containing 20-nt of the base-pairing sequence of the sgRNA and a 4-nt overhang complementary to BsmBI-digested pPEPZ-sgRNAclone (Figure 2C). The two oligos for each sgRNA were then annealed to produce duplex DNA with 4-nt overhang on both ends, and then the collection of 1499 sgRNA duplex DNAs were pooled together at equal molar concentrations, followed by phosphorylation. The phosphorylated sgRNA duplex DNA pool was then ligated into BsmBI-digested pPEPZ-sgRNAclone, resulting in the replacement of *mCherry* with *sgRNA* (Figure 2D). This ligation product could be directly transformed into *S. pneumoniae*. However, transformation into *E. coli* was preferred to allow visual red/white screening of cloning efficiency: colonies containing the parental (*mCherry*) vector are red, while colonies containing the *sgRNA* construct are white. In our study, no red colony showed up among the 70,000 colonies, indicating this method was extremely efficient in producing a high quality sgRNA pool. The *E. coli* library could also be used as a reservoir of sgRNAs for construction of reproducible CRISPRi libraries among *S. pneumoniae* D39V strains with different genetic backgrounds (e.g. mutant strains) or to transform other pneumococcal strains that contain the pPEPZ integration region, which is harbored by 64.21% of all sequenced pneumococcal strains (Keller et al., 2019). The plasmid pool was then transformed into *S. pneumoniae* D39V with the above described tet-inducible *dcas9* (Figure 2E). To compare the tet-inducible CRISPRi system to the IPTG-inducible CRISPRi system previously published by our group (Liu et al., 2017), the sgRNA library was simultaneously transformed into strain DCI23, which is *S. pneumoniae* D39V with IPTG-inducible *dcas9* (Liu et al., 2017).

### Benchmarking CRISPRi-seq with both tet- and IPTG-inducible libraries *in vitro*

CRISPRi screens with the constructed tet- and IPTG-inducible libraries were performed in C+Y medium, a standard laboratory pneumococcal growth medium (Figure 3A). Libraries were induced for approximately 21 generations with doxycycline and IPTG, respectively. The key Illumina elements, read 1 and read 2, inserted upstream and downstream of the sgRNA transcription unit, provided the binding sequence for the PCR handle (Figure 3A). With this design, the sgRNAs in the CRISPRi library could be amplified by a one-step PCR and sgRNAs subsequently quantified by Illumina sequencing. Hence, this process was named CRISPRi-seq. Sequencing verified the presence of all 1499 sgRNAs in the uninduced samples of both tet- and IPTG-inducible libraries. The relative abundance of sgRNAs ranged from 0.06 to 2.4 (Figure 3B), confirming that both CRISPRi libraries contain all designed sgRNAs. In uninduced samples, no bias towards any of the sgRNAs was detected, as their counts appeared to be normally distributed (Figure 3B). In addition, the sgRNA abundance in the two libraries was highly correlated (Figure S1A), confirming that our cloning strategy enabled repeatable transplantation of the sgRNA pool among parent strains with different genetic backgrounds. Induction of dCas9 (CRISPRi ON) by either doxycycline or IPTG resulted in a similar change in sgRNA profile (Figure 3B). The evaluated fitness, defined as the log2 fold change of sgRNA abundance upon induction, was highly consistent between the tet- and IPTG-inducible CRISPRi libraries (Figure 3C); only five sgRNAs exhibited a statistically different abundance (log2FC>1, *p*_adj_<0.05) (Figure S1B). The sgRNAs that were significantly less abundant upon dCas9 induction were categorized as targeting essential operons or genes. Likewise, sgRNAs that increased in abundance upon induction were defined as targeting costly operons or genes, while sgRNAs that did not change in abundance were defined as neutral. Based on this definition, 339 sgRNAs were defined as targeting essential operons or genes, 1160 sgRNAs defined as neutral, and none defined as costly for *S. pneumoniae* growth *in vitro* (supplementary table S3). Out of the 1499 sgRNAs, there were 1186 sgRNAs targeting individual genes, 162 sgRNAs targeting two-gene operons, and 151 sgRNAs targeting operons with three or more genes. Among these, 248 single-gene, 52 two-gene and 39 three-or-more-gene operons were found to be essential (Figure 3D). The majority of the essential genes defined by our CRISPRi-seq results have been previously identified as essential or responsive by Tn-seq studies (Liu et al., 2017; Opijnen and Camilli, 2012; van Opijnen et al., 2009), indicating high consistency between the approaches (Figure S2) (supplementary table S4).

**Figure 3.**
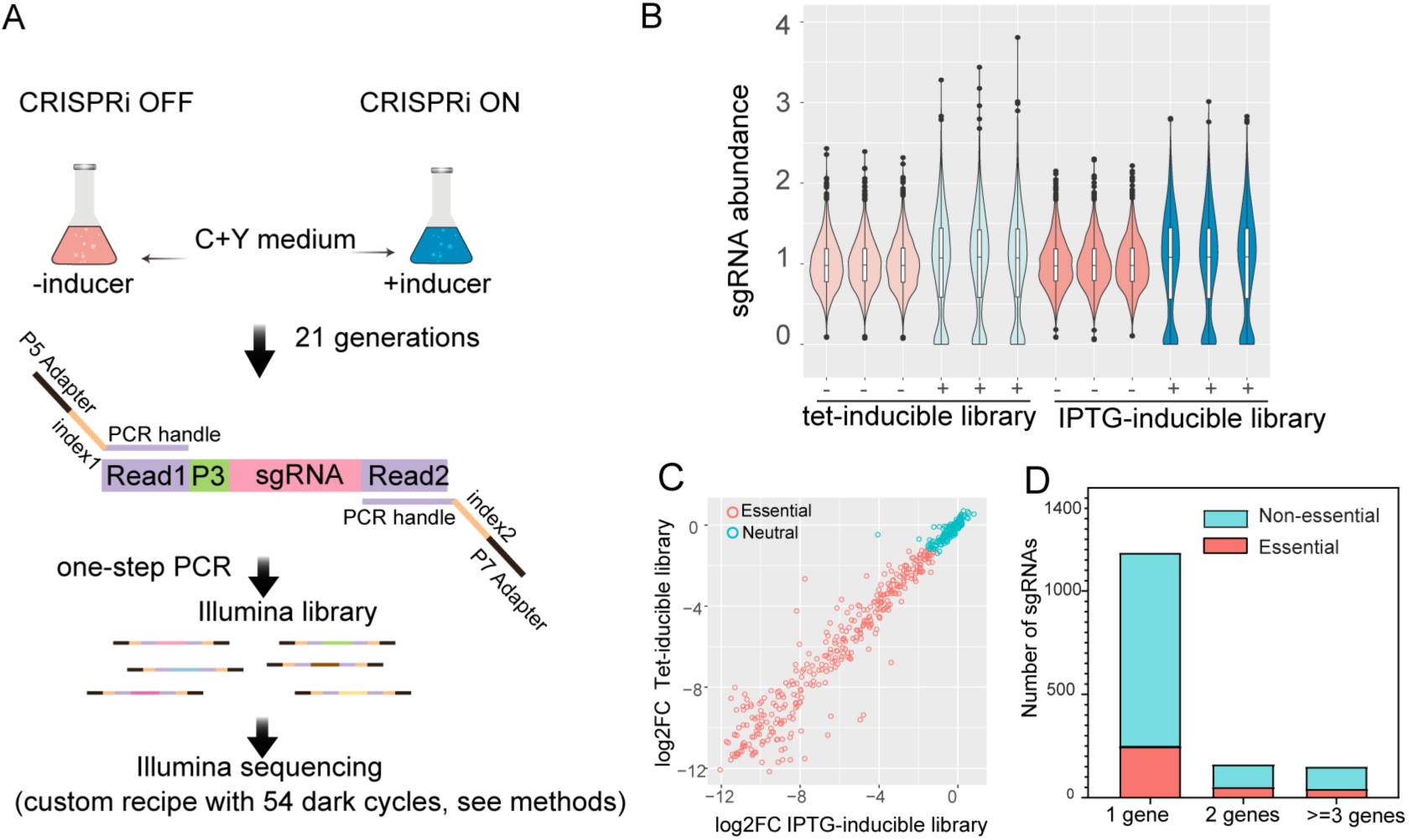
Fitness evaluation of CRISPRi targets under laboratory conditions. (A) Workflow of CRISPRi-seq. The tet- or IPTG-inducible CRISPRi libraries were cultured in C+Y medium in the absence (CRISPRi-OFF) or in the presence (CRISPRi-ON) of 10 ng/µl doxycycline or 1 mM IPTG. Bacteria were collected after approximately 21 generations of growth. Genomic DNA was isolated and used as a template for PCR. The forward oligo binds to Illumina amplicon element read 1 and contains the Illumina P5 adapter sequence; the reverse oligo binds to read 2 and contains the P7 adapter. Index 1 and index 2 were incorporated into the forward and reverse oligos respectively, for barcoding of different samples (see supplementary table S9). (B) Violin plots showing the distribution of sgRNA abundance in each samples. ‘-’ represents control samples without inducer; ‘+’ represents induced samples. The abundance of sgRNA =1499*(counts of sgRNA)/(total counts of all sgRNAs). (C) Correlation of the fitness of targets evaluated by IPTG-inducible and tet-inducible libraries. The log2FC, calculated with DEseq2, represents the fold change of sgRNA frequency between the control sample and induced sample. (D) Refinement of essential and non-essential genes of *S. pneumoniae* D39V by CRISPRi-seq. The sgRNAs were classified according to the number of their targets. 1 gene represents the sgRNAs targeting single gene operons; 2 genes represents the sgRNAs targeting two gene operons; >=3 genes represents the sgRNAs targeting operons with three or more genes.

### Bottlenecks and heterogeneity of *S. pneumoniae* in mouse pneumonia

During early infection, due to general stresses placed upon the bacterium within the host, such as nutrient restriction or innate immune system responses, a random part of the bacterial population might die off. This phenomenon is called a bottleneck, and the corresponding bottleneck size is the effective population size that gives rise to the final, post-bottleneck bacterial population that causes the infection. Such bottlenecks have been reported for pneumococcal infection and transmission (Gerlini et al., 2014; Kono et al., 2016; Opijnen and Camilli, 2012), but precise estimations of their sizes are lacking due to prior ineffective methodologies. A potential bottleneck in our pneumonia mouse model could cause random sgRNA loss from the pool in the CRISPRi-seq screens, instead of depletion through biological selection. This would introduce bias, in the sense that sgRNA depletion would no longer be an accurate proxy for gene essentiality. However, our set-up does allow for assessing both the presence and size of such a bottleneck, since we can accurately quantify the abundance of all 1499 strains in mice by Illumina sequencing, without inducing the CRISPRi system. Any loss of sgRNAs should then be attributable to a bottleneck effect, whose size was previously estimated on the basis of allele (here: sgRNA) frequencies in the pool before and after infection (Abel et al., 2015b).

To examine pneumococcal bottlenecks during a model of pneumonia, standard inbred adult BALB/c mice were infected with the *S. pneumoniae* CRISPRi library via the intratracheal route and CRISPRi-seq was performed from bacteria isolated at 48 hpi (from both lung and blood samples) and 24 hpi (only lung samples, as there are no detectable bacteria in the bloodstream at this time point) (Figure 4A). At 24 hpi, bottleneck sizes were relatively large, and covered more than 10-fold the library diversity in all samples except two (Figure 4B). Furthermore, estimated bottleneck sizes appeared to be smaller for CRISPRi-induced samples, which is likely due to the early drop-out of essential operons (Figure 4B).

**Figure 4.**
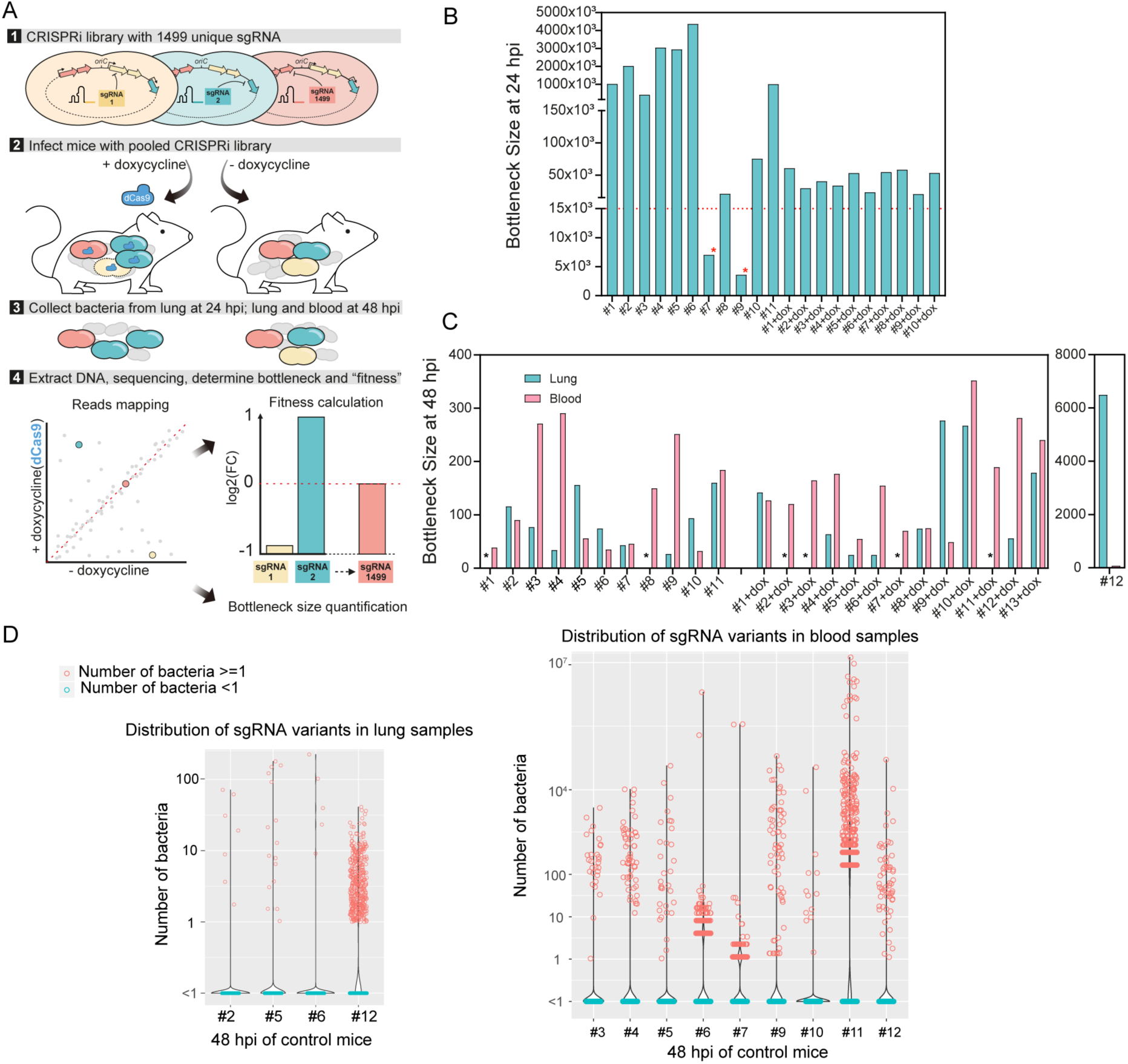
Exploring bottlenecks and pathogenesis of *S. pneumoniae* with CRISPRi-seq. (A) Workflow of fitness cost and bottleneck evaluation in a mouse pneumonia model by CRISPRi-seq. (B) Bottleneck size of lung samples at 24 hpi. 11 mice were treated with control chow, and 10 mice were treated with doxycycline chow. The horizontal red dash line marks 14,990 bacterial cells, which is a 10-fold theoretical coverage of the CRISPRi library. The red asterisks point to mouse #7 and mouse #9 in the control group. The bottleneck size of these two mice is lower than 10-fold of the library diversity. (C) Bottleneck size in lung and blood at 48 hpi. The black asterisks point out the lung samples without successful collection of bacterial samples, which include mice without doxycycline treatment #1 and #8, mice treated with doxycycline #2-dox, #3-dox, #7-dox, and #11-dox. (D) The number of bacteria barcoded with different sgRNAs in the control group (no doxycycline treatment) was calculated according to the bacterial load and sgRNA abundance in the population. Violin plots show the distribution of bacteria number in the lung samples (left panel) and blood samples (right panel), each dot represents one bacterial variant. Notice that some mice were not shown here, because the total bacterial load was below the limit of detection and the bacteria number of each variant cannot be calculated.

At 48 hpi, we observed a strong population size reduction in both lung and blood samples, and the bottleneck outcome was estimated to be as low as 25 bacterial cells responsible for causing disease (Figure 4C). CRISPRi induction did not seem to have a substantial effect on bottleneck size estimations, suggesting that the bottleneck selection effect overshadows the CRISPRi selection effect (Figure 4C). Surprisingly, bottleneck sizes varied considerably between replicates and did not correlate between lung and blood samples of the same host (Figure 4C). Moreover, there was little to no overlap in the different surviving strains in blood and lung samples within mice, indicating independent bacterial survival in lung and blood invasion (Figure S3). Taken together, these observations highlight the impact of bottlenecks on the outcomes of infection and strongly suggest that bacterial survival during infection in the mouse pneumonia model is highly heterogenous and bacterial survival is a stochastic event.

Quantification of the abundance of each mutant can provide information about bacterial replication and population expansion. To this end, we estimated the cell number of each mutant based on the abundance of each sgRNA in the library and the bacterial load in both lung and blood of the mice on control feed (Figure 4D). Dramatic stochastic changes in the genetic composition of the CRISPRi population were observed in all mice on control feed for both lung and blood samples at 48 hpi (Figure 4D). In addition, there was no correlation of bacterial genetic composition among samples from different mice, as individual mice have different dominant isogenic mutants. Most strains have 0 sgRNA reads, indicating most bacteria were cleared from the lungs or failed to invade the bloodstream. Some lowly abundant strains appear to have managed to survive, but not to actively multiply in both host niches, with bacterial number estimates between 1-10. However, especially in the blood samples, some variants reached high cell numbers (up to 10^7^), suggesting that invasion by a few sgRNA strains was followed by rapid replication. High replication rates in blood were further supported by observed bacterial loads in blood that were much higher than the estimated bottleneck sizes (Figure S4). Lastly, mouse number 12 seems to be less competent in clearing bacteria from the lung as clearly more variants survived, further stressing the importance of individual mouse effects despite being an inbred mouse strain (Figure 4C, D). Notice that here we used a published population level doubling time estimate (Opijnen and Camilli, 2012) for calculations of the bottleneck size. However, as described in Figure 4D, we observed subpopulations with divergent behaviors, indicating high degrees of heterogeneity of bacterial growth in both lung and blood during infection. This brings challenges for accurate estimation of doubling time at the population level. Different destinies of pneumococcal cells in the mouse infection model may be explained by bacterial phenotypic diversity or host-response diversity (Kreibich and Hardt, 2015). It has been determined that individual bacteria may occupy different micro-environments and can thus be exposed to dramatically different stimuli (Davis et al., 2015), which may contribute a level of randomness for certain pneumococcal clones to survive in the host. In addition, a single mouse passage can augment the virulence of some strains (Briles et al., 1981), such that within host evolution for genetic adaptation may lead to the emergence of subpopulations with different fates.

### CRISPRi-seq screen at 24 hpi identifies PurA as essential in a mouse pneumonia model

At 48 hpi, the effect of CRISPRi selection is overshadowed by a dramatic stochastic loss of mutants in the population while passing through the bottleneck, and thus this timepoint cannot be used to evaluate the fitness of targets by CRISPRi-seq (Figure 4C). However, earlier at 24 hpi, all mice except control mice numbers 7 and 9 exhibit a bottleneck size greater than 10-fold of the diversity in the CRISPRi library. Additionally, the CRISPRi induction *in vivo* clearly caused extra stress to the population, as the bottleneck size of the doxycycline-treated mice was smaller than control mice (Figure 4B). We thus analyzed the fitness of target genes during lung infection based on the sequencing data obtained at 24 hpi, excluding control mice numbers 7 and 9 (supplementary table S5).

*In vivo* fitness was compared to growth in laboratory media to identify those genes that became either more or less essential during infection (Figure 5). There were 46 sgRNAs whose targets showed significantly differential fitness between *in vivo* and *in vitro* conditions (log2FC>1, p_adj_<0.05), including 31 sgRNAs whose targets were more essential *in vivo* and 15 sgRNAs whose targets were less essential *in vivo* (supplementary table S5). Seeking to identify novel virulence factors, we next focused on those genes that scored as more essential *in vivo*.

**Figure 5.**
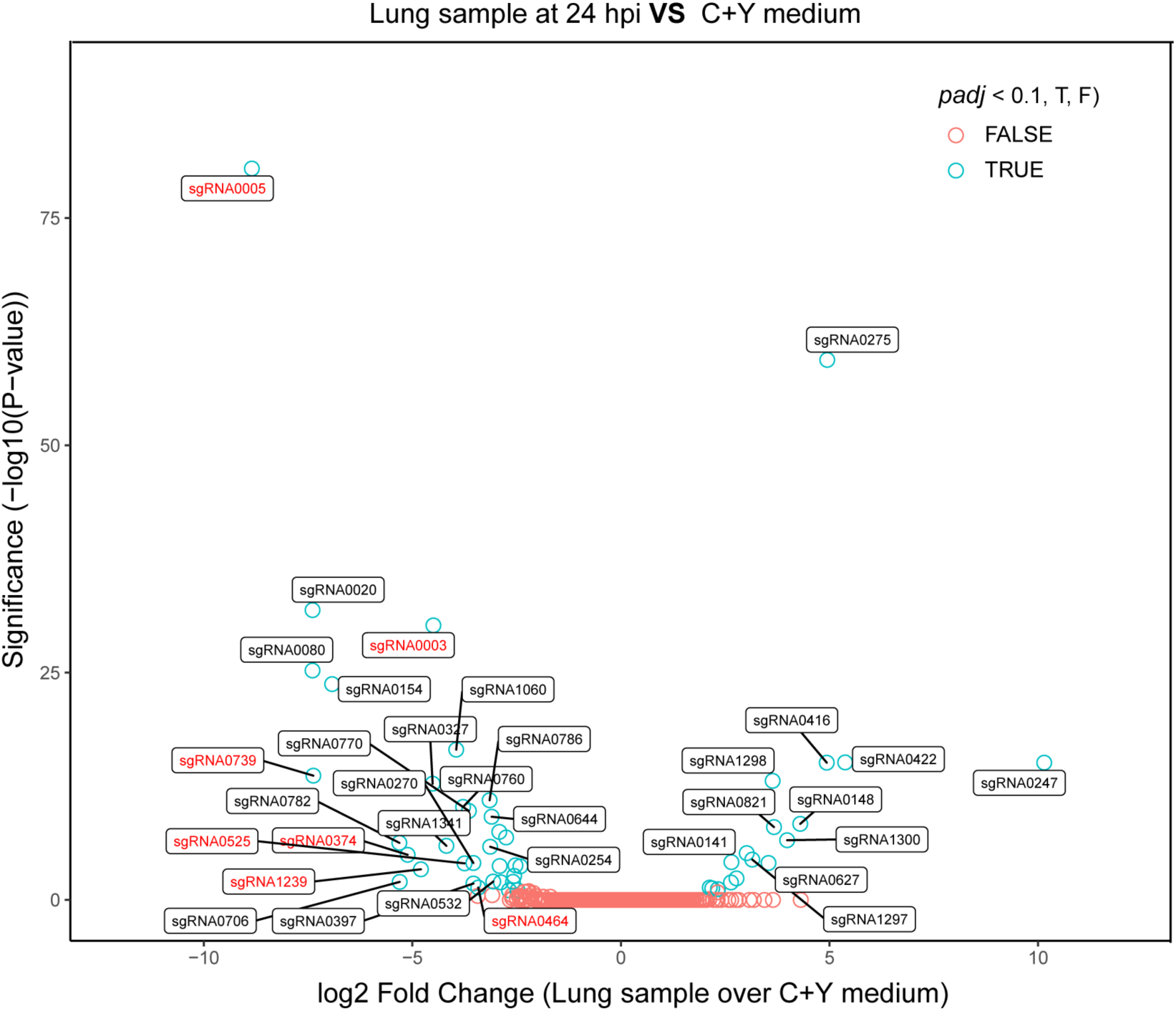
Comparison of fitness cost of gene depletion by CRISPRi by different sgRNAs between the mouse lung infection model at 24 hpi and laboratory C+Y medium. The difference was shown as the log2 fold change between the two conditions by DEseq2 analysis. The sgRNAs highlighted in red were the ones we selected for follow-up confirmation studies.

We selected 7 sgRNAs identified as targeting 13 neutral genes in C+Y laboratory growth media but predicted essential *in vivo* with a log2FC>3 difference (supplementary table S6). In line with the *in vitro* CRISPRi-seq data, most of the targeted genes (8) could be deleted without a detectable growth defect in C+Y medium (Figure 6A). Note that *spv_2285* was not tested, as it is encodes a degenerate gene (Slager et al., 2018). However, *divIC* (targeted by sgRNA0003), *spxA1* (sgRNA0464), and *dpr* (sgRNA0525) were identified as essential, since multiple attempts of deletion failed, corroborating the results of other studies (Liu et al., 2017; van Opijnen et al., 2009). Interestingly, for *pezT* and *pezA*, identified as an epsilon/zeta toxin-antitoxin system (Mutschler et al., 2011), single deletion of the toxin gene *pezT* or double deletion of *pezA-T* system were achieved and the resulting mutants showed no growth defect (Figure 6A). However, single deletion of the antitoxin gene *pezA* alone failed, an observation that indicates the *pezT-A* toxin-antitoxin system is active under *in vitro* growth conditions.

**Figure 6.**
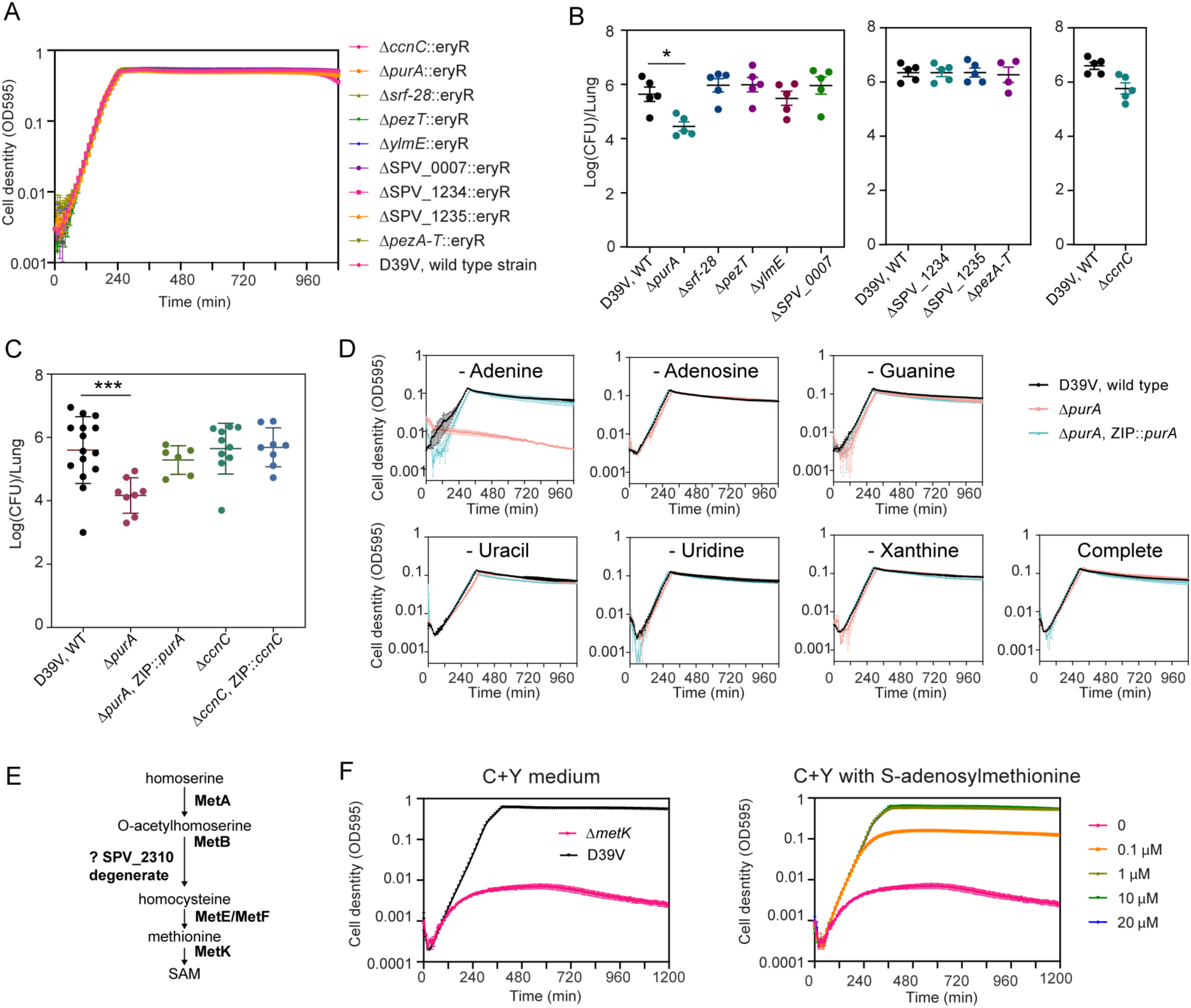
PurA is an important pneumococcal gene during an *in vivo* murine pneumonia model. (A) *In vitro* growth of the deletion mutants and the wild-type D39V strain in C+Y medium. Cell density was determined by measuring OD595nm every 10 minutes. The values represent averages of three replicates with SEM (same for panel D and E). (B-C) Mouse infection with individual mutants, compared to wild type D39V. Each dot represents a single mouse. Mean with SEM was plotted. (B) The mutants were tested in three batches of infection assays, for each assay the wild-type strain was tested in parallel. Significant difference between D39V and Δ*purA* was tested by Sidak’s multiple comparisons test, and the adjusted *p* value is 0.0158. Although the *ccnC* mutant showed slightly less bacterial counts in this experiment, this was not statistically significant. (C) Validation study of sgRNA0005 targets, *ccnC* and *purA*. The virulence of deletion mutants and complementary strains were tested and compared to wild type D39V. There was a significant difference between the wild-type and Δ*purA* strain tested by Dunnett’s multiple comparisons test, and the Adjusted *p* value is 0.0007. Note that ectopic expression of *purA* complemented the phenotype. (D) Growth of *purA* mutants in blood-like medium lacking adenine, adenosine, guanine, uracil, uridine, xanthine, and complete medium. (E) The pathway of S-adenosylmethionine (SAM) synthesis. (F) Growth of the *metK* deletion mutant in C+Y medium and C+Y medium supplemented with different concentrations of SAM.

Mice were then infected with the 8 viable knockout strains and the *pezA-T* double mutant individually by intratracheal challenge, and the bacterial load in the lung of each mutant compared to wild-type strain D39V (Figure 6B). Among the 9 mutants, *purA* (targeted by sgRNA0005), which showed the biggest log2FC between infection and C+Y medium in the CRISPRi-seq screen, was confirmed to be attenuated. sgRNA0005 targets an operon consisting of *ccnC* and *purA.* Infection experiments with knockout and complementary strains of these two genes confirmed that deletion of *purA*, but not *ccnC*, led to strongly attenuated *S. pneumoniae* virulence (Figure 6C). PurA, an adenylosuccinate synthetase, was previously identified to have a virulence role in experimental pneumococcal meningitis (Molzen et al., 2011). As adenylsuccinate synthetase is important for purine biosynthesis, we suspect the attenuated virulence of the *purA* knockout mutant is caused by lack of purine availability in the corresponding *in vivo* niches. To probe this further, we used a synthetic blood-like medium (BLM) (Aprianto et al., 2018) to propagate the *purA* mutants. The *purA* knockout mutant did not show a growth defect in BLM medium supplemented with complete nucleobases solution. However, specific reduction of adenine in the media led to marked growth retardation (Figure 6D).

### *In vitro* essential genes identified as non-essential *in vivo* highlight the power of CRISPRi-seq

Out of the 15 sgRNAs whose targets were identified to be significantly less essential *in vivo* than in laboratory C+Y medium, 11 sgRNAs were identified as essential in C+Y but neutral in lung infection (supplementary table S6). The Ami oligopeptide permease, involved in uptake of environmental nutrients (Claverys et al., 2000), is encoded by the *amiACDEF* operon. Interestingly, four sgRNAs targeting different genes of the Ami oligopeptide permease, *amiA*, *aimC*, *aimE*, and *amiF,* were independently identified to be neutral in lung infection but essential in C+Y medium. Consistent with these observations, Ami permease was previously reported to be conditionally essential and important for nasopharyngeal colonization but not for pneumococcal lung infection (Kerr et al., 2004).

sgRNA0247, targeting *metK*, showed the most significant difference between *in vitro* and *in v*ivo fitness among this class of sgRNAs. The *metK* gene encodes S-adenosylmethionine synthetase, which catalyzes the formation of S-adenosylmethionine (SAM) from methionine and ATP (Figure 6E). In line with its predicted function, the growth defect of the *metK* deletion mutant in C+Y medium could be rescued by addition of S-adenosylmethionine (SAM), with 1 µM SAM completely restoring wild-type growth rate (Figure 6F). In human serum, the SAM level was reported to be approximately 130 nM (Li et al., 2015). The nonessentiality of MetK in lung infection might be explained by the presence of SAM in host tissue. A previous study showed that *Mycobacterium tuberculosis* cannot scavenge intermediates of SAM and methionine biosynthesis from the host, and thus considered SAM- and methionine-related pathways as potential new drug targets (Berney et al., 2015). However, for *S. pneumoniae*, our study indicates that neither methionine nor SAM synthesis related pathway are essential for lung infection. Indeed, related sgRNAs, like sgRNA0530 targeting *metB*, and sgRNA0193 targeting *metEF* were also seen to be neutral in infection (supplementary table S5).

## Discussion

The principal contribution of this study is the development of a concise pooled CRISPR interference system, aided by the establishment of a new sgRNA assessment algorithm, suitable for high-throughput quantitative genetic interaction screening on a genome-wide scale for the important human pathogen *S. pneumoniae.* A main advantage of this concise doxycycline-inducible system is that it can be used for *in vivo* studies as the library size is small (1499 unique sgRNA) so bacterial loads can be low and sequence depth does not need to be high. Using this system, we were able to map infection bottlenecks in a murine model of pneumococcal pneumonia and show that as little as 25 individual bacterial cells can cause systemic disease. In addition, CRISPRi-seq reveals that there is a large within host and between host variability in dealing with pneumococcal infection, strongly suggesting that future work would benefit from a single cell analytical approach to study towards pneumococcal infection. It would be interesting to see which host immune response are most successful at increasing pathogen bottleneck sizes, and this information might inform on innovative therapies.

Genes identified as essential in laboratory medium but neutral in the host provide information to refine the list of new therapeutic targets for *S. pneumoniae*. For example, MetK, the SAM synthetase involved in SAM and methionine pathways, was previously identified as potential drug target for *M. tuberculosis* (Berney et al., 2015). In contrast, our study shows that MetK is not a promising target for pneumococcal disease since it is not essential *in vivo*. Another example includes FolD, which catalyzes the production of 10-formyltetrahydrofolate (10-formyl-THF). FolD was identified as neutral *in vivo* and essential in C+Y medium (supplementary table S5). The nonessentiality of FolD in the host may be explained by efficient production of 10-formyl-THF in mammalian cells (Ducker and Rabinowitz, 2017). In addition, our CRISPRi-seq screens showed that *metQ* and *metPN*, encoding the ABC transporter for methionine, were not essential for *S. pneumoniae* survival during pneumonia, in line with a previous study that used individual deletion mutants (Basavanna et al., 2013). Based on these corroborating observations, high-throughput *in vivo* evaluation of fitness cost of genes by CRISPRi-seq can provide useful information on the nutrient requirements during pneumococcal infection.

Out of the 7 sgRNAs that showed reduced *in vivo* fitness in the pooled CRISPRi-seq and were tested individually, only *purA* was confirmed. The CRISPRi-seq screen is a competitive assay, which can amplify the fitness defect of mutants. Some mutants that show attenuated virulence in competitive assays may have no such phenotype upon individual infection (Basavanna et al., 2009, 2013), which may explain why other candidates were not confirmed in the individual infection study. Alternatively, while at 24 hpi the sgRNA coverage was still greater than 10-fold that of the library, the bacterial population is aggressively being cleared by the host innate immune system. Thus the assay most strongly tests for mutants that are more or less resistance to host clearance rather than mutants that are attenuated in replication. Regardless of their etiology, this study shows the existence of large bottlenecks in the commonly studied pneumococcal mouse pneumonia model, such that from an initial inoculum of 10e7 bacteria, in some cases, only 25 single bacteria established systemic disease. This finding begs the question of how realistic this small animal system is for modeling human pneumococcal pneumonia. Future studies might apply the CRISPRi-seq tool to other established pneumococcal models of disease such as the influenza superinfection model, the zebrafish meningitis model and the *Galleria mellanonela* larvae invertebrate model (Cools et al., 2019; Jim et al., 2016; Rudd et al., 2016; Saralahti et al., 2014).

In summary, the here presented concise CRISPRi-seq setup can be used for studying pneumococcal pneumonia, including bottleneck exploration. The library, it’s design rules and the underlying bioinformatic approaches developed here can now be expanded to study other infection-relevant conditions including testing of wild-type and knockout mouse strains and evaluation of antibiotics and other therapeutic interventions and may serve as an example for studies on other host-microbe interactions including human pathogens.

## Materials and Methods

### Bacterial strains and growth medium

*Streptococcus pneumoniae* D39V (Slager et al., 2018) was used as the parent strain for this study. C+Y medium (pH=6.8), and Columbia agar plate supplied with 5% sheep blood were used to grow the strain and its derivatives. Working stock of the pneumococcal cells, named as “T2 cells”, were prepared by collecting cells at OD600 0.3 and then resuspending with fresh C+Y medium with 17% glycerol, and stored at -80°C. *Escherichia coli* stbl3 was used for subcloning of plasmids. LB agar with 100 µg/ml spectinomycin was used to select the *E. coli* transformants. Strains and plasmids used in this study are listed in supplementary table S7. The oligos used for construction of mutants and strains used in this study are listed and described in supplementary table S8.

### Construction of a tetracycline inducible (tet-inducible) CRISPRi system in S. pneumoniae D39V

The tetracycline inducible CRISPRi system was constructed based on our previously published IPTG-inducible CRISPRi system in *S. pneumoniae* (Liu et al., 2017) and a newly developed pneumococcal tet-inducible platform (Sorg et al., 2019). First, a constitutively expressed pneumococcal codon-optimized *tetR* driven by promoter PF6 was amplified from D-T-PEP9Ptet (Sorg et al., 2019) and integrated into the chromosome at the *prs1* locus in D39V strain, resulting in strain VL2210. Three fragments were assembled to make the Ptet-*dcas9* construct for integration at the *bgaA* locus. Fragment 1 containing upstream of *bgaA* and *tetM* was amplified from DCI23 (Liu et al., 2017) and digested with XbaI; fragment 2 containing tet-inducible promoter PT4-1, here named Ptet, was amplified from plasmid pPEP8T4-1 (Sorg et al., 2019), and digested with XbaI/NotI; fragment 3 containing the coding region of *dcas9* and downstream of *bgaA* locus was amplified from strain DCI23 and digested with NotI. The three fragments were then ligated followed by transformation into VL2210 by selecting with 1 µg/ml tetracycline, resulting in strain VL2212.

### Construction of the dual fluorescent reporter strain and confocal microscopy

The codon optimized mNeonGreen was digested from pASR110 (pPEPZ-Plac-mNeonGreen) with BglII and XhoI and cloned into pPEPY-Plac (Keller et al., 2019), followed by transformation into strain VL2212, resulting in strain VL2339. The DNA fragment for insertion of *hlpA*-*mScarlet-I* was amplified from strain VL1780 (Kurushima et al., 2020) and transformed into VL2339, resulting in the final dual fluorescent reporter strain VL2351. To verify doxycycline levels were sufficient *in vivo* to induce inhibition via CRISPRi, mice were switched to feed containing doxycycline (or control feed) two days prior to infection, and then infected with the reporter strain via intra-tracheal infection, during which time mice remained on doxycycline feed (or control feed). At 48 hours post infection, whole blood was collected via cardiac puncture followed by hypotonic lysis of red blood cells and subsequent resuspension of remaining cells in PBS. Samples were placed on a glass slide, heat fixed and mounted in Cytoseal. Slides were imaged using a Leica TCS SPE Confocal microscope with a 63X objective, LAS X acquisition software, and processed using FIJI (Schindelin et al., 2012).

### Construction of knockout and complementary mutants in S. pneumoniae

The erythromycin resistant marker, encoded by *eryR*, was used as selection marker for the knockout mutants. Three fragments was assembled by Golden Gate cloning with either BsaI or BsmBI for each knockout mutant: Fragment 1 containing upstream of the target gene including its promoter sequence; fragment 2 containing *eryR* coding region with RBS; fragment 3 containing downstream of the target gene. The assembled DNA fragment was then transformed into D39V with 0.5 µg/ml erythromycin for selection. Notice that for making the Δ*metK* strain, 10 µM SAM was supplemented in the agar plate. To make the complementary strains, the target gene with its native promoter was amplified from genomic DNA of D39V and ligated with upstream and downstream homologous fragments of ZIP locus followed by transformation into the knockout mutant with 100 µg/ml spectinomycin for selection. Primers used here are listed in supplementary table S8.

### Construction of the pooled CRISPRi library

#### Construction of vector pPEPZ-sgRNAclone for sgRNA cloning by Golden Gate cloning

Integration vector pPEPZ (Keller et al., 2019) was used as backbone. A gBlock containing Illumina read 1 sequence, P3 promoter, *mCherry* flanking with BsmBI sites, dCas9 handle binding and terminator region of sgRNA, Illumina read 2 sequence, 8 bp Illumina index sequence and P7 adaptor sequence in order, was synthesized by Integrated DNA Technologies (IDT). In this design, *mCherry* provides the sgRNA base-pairing cloning sites and will be replaced with 20 bp specific sequence for targeting different genes. The Illumina sequences across the sgRNA cloning sites work as primer binding handles for one-step PCR amplification of the sgRNA sequence, in order to prepare amplicon library for Illumina sequencing. The gBlock was digested with BamHI/XholI, and then ligated with BamHI/XholI digested pPEPZ, followed by transformation into *E. coli* stbl3 selected with 100 µg/ml spectinomycin. The *E. coli* strain with this vector forms bright red colonies. The vector pPEPZ-sgRNAclone was deposited at Addgene (catalog #141090).

#### Selection of sgRNAs for each operon

Primary operons (pTSS operons) were annotated in *S. pneumoniae* D39V strain in a previous study (Slager et al., 2018). First, for all the identified pTSS operons, one sgRNA with high specificity and close proximity to the pTSS was designed for each operon. For genes that are not covered by pTSS operons, one sgRNA was selected for each gene. *S. pneumoniae* has multiple types of repeat regions, such as BOX elements, Repeat Units of the Pneumococcus (RUP), SPRITEs and IS elements (Slager et al., 2018). There are some sgRNAs targeting genes located in repeat regions, and as such these sgRNAs have multiple targeting sites. The sgRNAs and targets are listed in supplementary table S1. Post-hoc target identification, including off-target sites, was performed with a custom R script (https://github.com/veeninglab/CRISPRi-seq), of which the results are shown in supplementary table S2 and analyzed separately (https://www.veeninglab.com/crispri-seq, “Pneumococcal sgRNA library efficiency exploration”).

#### Cloning of sgRNAs by Golden Gate cloning

Two oligos were designed for each sgRNA (Figure 2). The two oligos were then annealed in TEN buffer (10 mM Tris, 1 mM EDTA, 100 mM NaCl, pH 8) in a thermocycler, 95°C for 5 minutes followed by slowly cooling down to room temperature. The annealed oligos were then pooled together at equimolar concentration, followed by phosphorylation with T4 polynucleotide kinase (New England Biolabs). The vector pPEPZ-sgRNAclone was digested with BsmBI and carefully purified by gel extraction to ensure removal of the *mCherry* fragment. The annealed oligos and digested pPEPZ-sgRNAclone were then ligated with T4 ligase, followed by transformation into *E. coli* stbl3 and selected with 100 µg/ml spectinomycin on LB agar plates. In total, more than 70,000 individual transformant colonies were obtained and collected, providing about a 50 fold theoretical coverage of the 1499 sgRNAs. No red colonies were visually present, indicating a very low false positive rate of the cloning. The oligos for cloning of sgRNAs are listed in supplementary table S8.

#### Construction of the pooled CRISPRi library in S. pneumoniae D39V

Plasmids from the *E. coli* library with the sgRNA pool were isolated, and transformed into *S. pneumoniae* VL2212 (this study) and DCI23 (Liu et al., 2017) to construct tet-inducible and IPTG-inducible CRISPRi-seq libraries, respectively. More than 10^7^ individual transformant colonies were obtained and collected for both of the strains.

#### CRISPRi-seq screen in laboratory medium

The screen was done over about 21 generations of growth in triplicates. The pooled libraries were grown in C+Y medium at 37ºC to OD_595_=0.3 as preculture. Then, the precultures were diluted 1:100 into C+Y medium with or without inducer, 1 mM IPTG or 10 ng/ml doxycycline. When OD_595_ reached 0.3, cultures were diluted into fresh medium by 1:100 again. Another 1:100 dilution was done in the same fashion, so in total three times of 1:100 dilution, ensuring about 21 generations of induction and competition (doubling time of approximately 26 min). Bacteria were collected when OD_595_=0.3 after the third dilution, and the pellets were used for gDNA isolation with the Wizard Genomic DNA Purification Kit (Promega) as described previously (Liu et al., 2017). The fitness evaluated by IPTG- or tet-inducible library is listed in supplementary table S3.

#### CRISPRi-seq screen in a mouse pneumonia model

The UCSD Institutional Animal Care and Use Committee approved all animal use and procedures (protocol number S00227M, Victor Nizet). Two days prior to infection, 6-8 week old female BALB/c mice (Jackson Laboratories - 000651) were fed control feed or 200 ppm doxycycline feed *ad libitum* (Envigo TD.120769, with blue food coloring), allowing serum concentrations of doxycycline to stabilize prior to infection (Redelsperger et al., 2016). Bacterial libraries were grown *in vitro* in C+Y medium in the absence of selection (i.e. no doxycycline) to an OD_600_ of 0.4, sonicated for 3 seconds to break up clumps, and then resuspended in PBS at a concentration of 1 × 10^8^ CFU per 30 µL. Mice were anesthetized with 100 mg/kg ketamine and 10 mg/kg xylazine (intraperitoneal administration), vocal cords were visualized with an otoscope and 30 µL bacteria was delivered into the lungs by pipetting. Mice were returned to the same cages after infection, containing doxycycline or control feed. At 24 or 48 hpi, mice were euthanized via CO_2_ asphyxiation, lungs were dissected and homogenized in 1 mL PBS, while blood was collected by cardiac puncture in the presence of EDTA to prevent clotting. Following tissue harvest, lung homogenate or blood was diluted in 15 mL C+Y medium (without selection), incubated at 37 °C with 5% CO_2_ until cultures reached an OD_600_ of 0.4. Samples were then pelleted and frozen before subsequent gDNA isolation and sequencing. For the comparison of fitness between mice infection model and C+Y medium, please refer to supplementary table S5.

#### sgRNA library target and efficiency evaluation

All potential sgRNA binding sites on the *S. pneumoniae* D39V genome were identified using the R package CRISPRseek (Zhu et al., 2014), taking into account PAM presence and allowing up to eight mismatches between spacers and genome. We set the maximum number of allowed mismatches to eight, because of (1) the exponential growth of computation time with this parameter and (2) any potential effect on a site with >8 mismatches was assumed to be negligible. The *S. pneumoniae* D39V genome (Slager et al., 2018) was downloaded from NCBI (CP027540.1) and read into R using the biomartr package (Drost and Paszkowski, 2017). All identified binding sites can be found in supplementary table S2.

In addition to the standard CRISPRseek output, we assessed for each binding site if it overlapped with any genetic element annotated with a *locus_tag* key in the GFF file on the non-template (NT) strand. If any, the locus tag was added to the table (“NTgene”), as well as which part of the sgRNA corresponding to that binding site was overlapping (“coverPart”: complete, 5′- or 3′-end) and with how many base pairs (“coverSize”) including the PAM. In case one binding site overlapped multiple annotated elements, both were inserted as a row in the table, with matching “site” numbers.

Furthermore, we estimated the relative retained repression activity (“reprAct”) of each sgRNA binding site compared to a hypothetical zero-mismatch binding site on the same locus, based on the mismatches with the sgRNA spacer. Retained repression activity depends on both the number and the within-spacer position of mismatches (Qi et al., 2013). Furthermore, the retained activity for an sgRNA with two adjacent mismatches appears to be the product of their individual retained scores, relative to a zero-mismatch silencing effect (Qi et al., 2013). We assumed this multiplication principle also holds for >2 and non-adjacent mismatches. Therefore, we computed per sgRNA, per binding site, the expected repression activity as the product of the nucleotide-specific retained activity scores as reported by Qi et al. (2013), estimated from their Figure 5D and averaged over the three intra-spacer regions they defined (Qi et al., 2013). The resulting score represents the estimated retained repression activity of the sgRNA on the binding site, relative to an hypothetical binding site for the same sgRNA on the same chromosomal locus, on the [0,1] interval.

According to this method, the maximum repression effect of any site with >8 mismatches would be 0.77% of the hypothetical zero-mismatch effect. This was indeed considered negligible, supporting our decision to only consider sgRNA binding sites with eight mismatches or less.

Lastly, we added to the table the relative distance of the binding site to the start codon of the genetic element it binds to, if any (“dist2SC”). This distance is normalized to the [0,1] interval using feature scaling, where a distance of 0 means binding on the start codon or partially overlap with the 5’-end of the element, and a distance of 1 means binding at or partial overlap with the far 3’-end of the element. Smaller distances are associated with more efficient transcription repression (Qi et al., 2013).

The custom R script used to produce this table can be found on Github (https://github.com/veeninglab/CRISPRi-seq). The script is written in a generic way, allowing to run the complete pipeline described above for any given NCBI genome, as we did for *S. pneumoniae* strains TIGR4 (AE005672.3), R6 (AE007317.1), Hungary19A-6 (CP000936.1), Taiwan19F-14 (CP000921.1), 11A (CP018838.1) and G54 (CP001015.1). Results tables and analysis of these genomes can be found on the Veeninglab website (https://www.veeninglab.com/crispri-seq).

#### Library preparation, sequencing and data analysis

The Illumina libraries were prepared by one-step PCR with oligos listed in supplementary table S8. The isolated gDNAs of *S. pneumoniae* were used as template for PCR. The index 1, index 2 and adapter sequence were introduced by this one-step PCR. For each 50 µl of PCR reaction, 4 µg of gDNA was used as input template, which enables us to obtain sufficient PCR products with as little as 8 cycles of PCR. In addition, we have tested 10 cycles, 20 cycles and 30 cycles of PCR reaction, and no significant difference was observed (data not shown), indicating no detectable bias introduced by PCR. The amplicons (304 bp) were then purified from a 2% agarose gel. Concentrations of amplicons were then determined by a Qubit assay (Q32854, ThermoFisher Scientific). Purified amplicons were sequenced on a MiniSeq (Illumina) with a custom sequencing protocol. The first 54 cycles for sequencing of common sequence of amplicons were set as dark cycles, and the following 20 cycles were used for sequencing of the diversified base-pairing region of sgRNA. The fastq files generated from sequencing are uploaded to the Sequence Read Archive (SRA) on NCBI with accession number PRJNA611488.

The 20 bp base-pairing sequences were trimmed out from read 1 according to their position with Trimmomatic Version 0.36 (Bolger et al., 2014). To map the sgRNA sequences, a pseudogenome containing all the sgRNA sequences was prepared, and the sgRNA sequences on the pseudogenome were annotated with sgRNA numbers, 1 to 1499. Then the trimmed reads were mapped to the pseudogenome with Bowtie 2 (Langmead and Salzberg, 2012). The sgRNAs were counted with featureCounts (Liao et al., 2014). The count data of sgRNAs were then analyzed with the DESeq2 package in R for evaluation of fitness cost of each sgRNA. We tested against a log2FC of 1, with an alpha of 0.05. Whenever log2FC are visualized or reported, these are shrunk with the apeglm method. The R script used for analysis is available at https://github.com/veeninglab/CRISPRi-seq. The size of infection bottlenecks was calculated as reported previously (Abel et al., 2015). The doubling time of *S. pneumoniae* used in the calculation was based on a previous Tn-seq study (Opijnen and Camilli, 2012) as 108 minutes.

#### Growth assays and luciferase assay

For Figure 1B, 6A and 6F, the working stock of each mutant, T2 cells, were thawed and diluted 1:100 into fresh C+Y medium, or C+Y medium with doxycycline at different final concentrations, or with different concentrations of S-(5’-Adenosyl)-L-methionine (A7007, Sigma Aldrich), as the initial cell culture. For Figure 6D, the T2 cells were thawed and diluted 1:100 into fresh Blood Like Medium (BLM) without nucleobases solution, or supplemented with individual nucleobases component (adenine, adenosine, guanine, uracil, uridine and xanthine), or with all the components (Aprianto et al., 2018). Then 300 µl of the initial culture was aliquoted into each well of 96-well plates with 3 replicates. Cell density were monitored by measuring OD595 every 10 minutes with a Tecan Spark microtiter plate reader at 37ºC. Luciferase assay (Figure 1B) was performed as previously described (Liu et al., 2017). Luciferin (D-Luciferin sodium salt, SYNCHEM OHG) was added into C+Y medium at final concentration of 450 µg/ml as substrate of the luciferase. Luminescence signal were measured every 10 minutes with a Tecan Spark microtiter plate. The growth curve and luciferase activity curve were plotted with GraphPad Prism 8 as described previously (Sorg and Veening, 2015).

## Author Contributions and Notes

X.L, J.M.K and J.W.V designed research, X.L. and J.M.K performed research, V.D.B. wrote the R script for sgRNA evaluation and DEseq2 analysis, X.L, J.M.K, V.D.B, V.N and J.W.V analyzed data; and X.L, J.M.K, V.D.B, V.N and J.W.V wrote the paper. The authors declare no conflict of interest. This article contains supporting information online.

### Acknowledgments

We are grateful to Lance Keller for critical reading of the manuscript. We appreciate all members of the Veening lab for stimulating discussions. This work was supported by the Swiss National Science Foundation (SNSF) (project grant 31003A_172861 to J.W.V.) and JPIAMR grant (40AR40_185533 to J.W.V.) from SNSF. Work in the Nizet lab is supported by NIH grant AI145325. J.M.K. was supported by the University of California President’s Postdoctoral Fellowship Program (UC PPFP).

