## Supplementary figures and Tables S6, S7 and S9 for "Exploration of bacterial bottlenecks and *Streptococcus pneumoniae* pathogenesis by CRISPRi-seq"

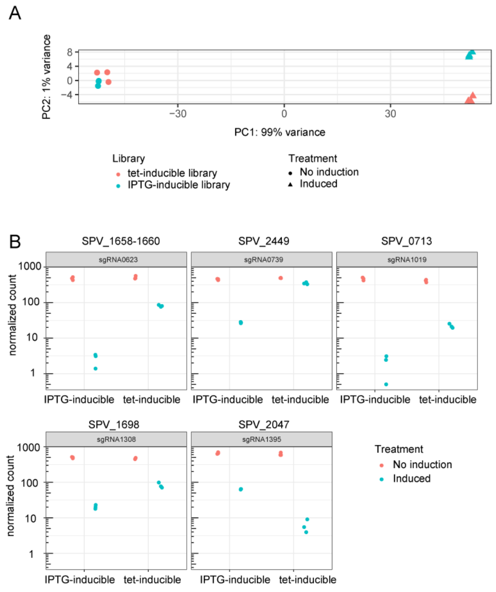


**Figure S1.** (A) PCA analysis of the samples grown in C+Y medium with or without inducer. (B) The sgRNAs that showed significantly different fold change of control sample and induced samples between the tet-inducible library and IPTG-inducible library

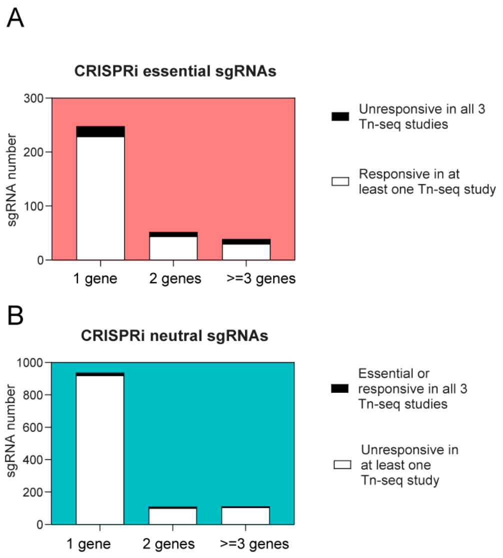


**Figure S2.** Comparison of the essential gene list identified by CRISPRi-seq and Tn-seq.

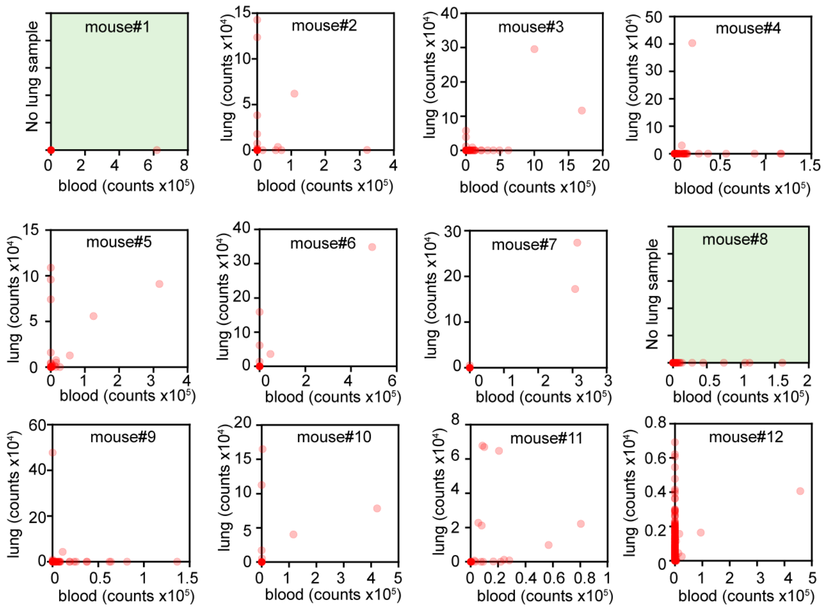


**Figure S3.** Correlation of sgRNA abundance between lung and blood samples per mouse at 48 hpi, in the control group (not treated with doxycycline). Note for mouse #3 and mouse #8, we failed to collect bacteria from lung samples, so only the sgRNA abundance of blood samples was shown for these two mice.

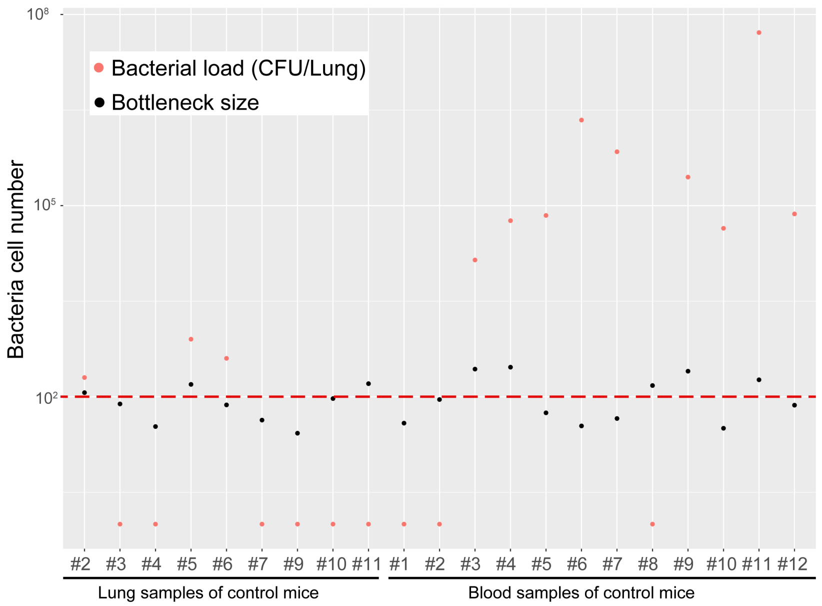


**Figure S4.** Comparison of bacterial load to estimated bottleneck size. Bacterial load (CFU/Lung) was determined by plating homogenized tissue and numerating colony numbers. Notice that 100 CFU/Lung is the detective limits (red dash line). The red dots below 100 represents bacterial load under detective limits. Bottleneck size was estimated on the basis of allele (here: sgRNA) frequencies in the pool before and after infection as described in the materials and methods.

**Supplementary Tables:**

**Supplementary Tables S1-S5 and S8 are provided as separate excel files.**

**Table S6**. sgRNAs that showed significant differential essentiality between the C+Y medium and the infection model.

| sgRNA | C+Y medium | Pneumonia | Interaction (log2FC) | Targets |
|---|---|---|---|---|
| **sgRNA0003** | neutral | essential | -4.43 | SPV_0007, *divIC* (SPV_0008) |
| **sgRNA0005** | neutral | essential | -8.76 | *purA* (SPV_0024), *ccnC* (SPV_2078) |
| **sgRNA0374** | neutral | essential | -4.67 | SPV_2285, *pezT*(SPV_0931), *pezA*(SPV_0930) |
| **sgRNA0464** | neutral | essential | -3.00 | SPV_1234, SPV_1235, *spxA1*(SPV_1236) |
| **sgRNA0525** | neutral | essential | -3.49 | *dpr* (SPV_1402) |
| **sgRNA0739** | neutral | essential | -7.07 | *srf-28* (SPV_2449) |
| **sgRNA1239** | neutral | essential | -4.25 | *ylmE* (SPV_1478) |
| sgRNA0247 | essential | neutral | 9.54 | *metK* (SPV_0664) |
| sgRNA0275 | essential | neutral | 4.91 | *folD* (SPV_0721) |
| sgRNA0416 | essential | neutral | 4.79 | *fhs* (SPV_1087) |
| sgRNA0148 | essential | neutral | 4.10 | SPV0328-0390 |
| sgRNA1300 | essential | neutral | 3.78 | *amiC* (SPV_1670) |
| sgRNA1298 | essential | neutral | 3.55 | *amiE* (SPV_1668) |
| sgRNA0627 | essential | neutral | 3.32 | *amiA* (SPV_1671) |
| sgRNA1297 | essential | neutral | 3.01 | *amiF* (SPV_1667) |
| sgRNA0141 | essential | neutral | 2.91 | *glyP* (SPV_0372) |
| sgRNA1121 | essential | neutral | 2.63 | *recN* (SPV_1062) |
| sgRNA0683 | essential | neutral | 2.57 | *ulaH* (SPV_1839), *yajC* (SPV_1838) |

**Notes:**

**sgRNA** shows the sgRNA number in the library, sgRNAs in bold were the 7 sgRNAs that we selected for follow-up confirmation study; **C+Y medium** and **Pneumonia** show the categories of the sgRNAs based on analysis of DEseq2 with the fitness of targets in C+Y medium and in lung of the pneumonia model at 24 hpi, respectively. **Interaction (log2FC)** represents the difference of sgRNA frequency fold change of the control sample and induced sample between C+Y medium and pneumonia model. **Targets** are the genes in the targeted operon of the sgRNAs.

**Table S7. Strains and plasmids used in this study**

| Strains/Plasmids | Genotype | Reference |
|---|---|---|
| *S. pneumoniae* | | |
| D39V | Serotype 2 strain, wild-type | (Slager et al., 2018) |
| DCI23 | D39V, Δ*bgaA*::*$P_{lac}$-*dcas9sp* (tet$^R$); Δ*prs1*::PF6-*lacI* (Gm$^R$ ) | (Liu et al., 2017) |
| VL1780 | D39V, *hlpA*::*hlpA_hlpA-mScarlet-I* (cam$^R$) | (Kurushima et al., 2020) |
| XL28 | D39V, Δ*bgaA*::*$P_{lac}$-*dcas9sp* (tet$^R$); Δ*prs1*::PF6-*lacI* (Gm$^R$ ); **cil*::P3-*luc* (kan$^R$), Δ*CEP*::P3-*sgRNAluc* (spec$^R$) | (Liu et al., 2017) |
| D-T-PEP9Ptet | D39V, Δ*prs1*::PF6-*tetR* (Gm$^R$ ); Δ*CEP*::$P_{tet}$-*luc-gfp* (spec$^R$) | (Sorg et al., 2019) |
| VL2210 | D39, Δ*prs1*::PF6-*tetR* | This study |
| VL2212 | D39, Δ*prs1*::PF6-*tetR*, Δ*bgaA*:: *$P_{tet}$-*dcas9* | This study |
| VL2339 | D39, Δ*prs1*::PF6-*tetR*, Δ*bgaA*:: *$P_{tet}$-*dcas9*, *cil**::$P_{lac}$-*mNeonGreen* (Kan$^R$) | This study |
| VL2351 | D39, Δ*prs1*::PF6-*tetR*, Δ*bgaA*::$P_{tet}$-*dcas9*, *cil**::$P_{lac}$-*mNeonGreen* (Kan$^R$), *hlpA*::*hlpA-mScarlet-I* (cam$^R$) | This study |
| VL3106 | D39V, Δ*ccnC*::eryR | This study |
| VL3107 | D39V, Δ*purA*::eryR | This study |
| VL3108 | D39V, Δ*srf-28*::eryR | This study |
| VL3109 | D39V, Δ*pezT*::eryR | This study |
| VL3110 | D39V, Δ*ylmE*::eryR | This study |
| VL3111 | D39V, ΔSPV_0007::eryR | This study |
| VL3112 | D39V, ΔSPV_1234::eryR | This study |
| VL3113 | D39V, ΔSPV_1235::eryR | This study |
| VL3114 | D39V, Δ*pezA-T*::eryR | This study |
| VL3168 | D39V, Δ*ccnC*::eryR, *ZIP**::P-*ccnC* (Native promoter of *purA-ccnC* operon was used) | This study |
| VL3169 | D39V, Δ*purA*::eryR, *ZIP**::P-*purA* (Native promoter of *purA-ccnC* operon was used) | This study |
| VL3462 | D39V, Δ*metK*::eryR | This study |
| Plasmids | | |
| pPEPZ-sgRNAclone | *ZIP**', specR, P3, mCherry, sgRNA(dCas9handling+terminator), '*ZIP** | This study |
| pPEP8T4-1 | *CEP**', spec$^R$, *tetR*, PT4-1, *luc-gfp*, '*CEP** | (Sorg et al., 2019) |
| pASR110 (pPEPZ-$P_{lac}$-mNeonGreen) | *ZIP**', spec$^R$, $P_{lac}$-mNeonGreen, '*ZIP** | (Keller et al., 2019) |
| pPEPY-$P_{lac}$ | *cil**', kan$^R$, $P_{lac}$, MCS, '*cil** | (Keller et al., 2019) |
| pJWV502 | *ampR, bgaA', tet$^R$,* $P_{Zn}$-*gfp, ery$^R$, 'bgaA* | (Liu et al., 2017) |

Notes:

1. $cam^R$: chloramphenicol resistance; $ery^R$: erythromycin resistance; $Gm^R$: gentamycin resistance; $kan^R$: kanamycin resistance; $spec^R$: spectinomycin resistance; $tet^R$: tetracycline resistance.
2. *$P_{lac}$: The IPTG inducible promoter
3. *$P_{tet}$: the tetracycline inducible promoter. The here used $P_{tet}$ represents PT4-1in our previous study (Robin's)
4. *cil: chromosome intergration locus, represents a non-coding region between SPD_0422 and SPD_0423.
5. *ZIP:

Sequence (5'-3') of the gBlock containing Illumina read 1 sequence, P3 promoter, *mCherry* flanking with BsmBI sites, dCas9 handle binding and terminator region of sgRNA, Illumina read 2 sequence, 8 bp Illumina index sequence and P7 adaptor sequence in order

GATCTAGCAGATCTGAGAGGATCCCCATTCTACAGTTTATTCTTGACATTGCACTGTCCCCCTGGTATA
ATAACTATATGAGACGAGGAGGAAAATTAATGAGCAAAGGAGAAGAAGATAACATGGCAATCATCAA
AGAATTTATGCGTTTCAAAGTTCACATGGAAGGTTCTGTAAACGGACACGAATTTGAAATTGAAGGTG
AAGGTGAAGGCCGTCCTTATGAAGGAACACAAACGGCAAAGCTGAAAGTAACAAAAGGCGGACCGCT
TCCGTTTGCATGGGATATCCTTTCTCCGCAATTCATGTACGGTTCAAAAGCATACGTGAAGCATCCGGC
TGATATTCCTGATTATTTGAAGCTGTCATTCCCTGAAGGCTTCAAATGGGAGCGTGTGATGAACTTTGA
AGATGGCGGTGTTGTTACTGTTACTCAAGATTCAAGCCTTCAAGACGGTGAATTTATTTACAAAGTGA
AGCTGCGCGGAACAAACTTCCCATCTGACGGACCTGTCATGCAAAAGAAAACAATGGGCTGGGAAGC
AAGCTCTGAACGCATGTATCCAGAGGACGGTGCTTTAAAAGGAGAAATCAAACAGCGTTTGAAGCTG
AAAGACGGCGGACACTATGACGCTGAAGTGAAAACAACTTACAAAGCGAAAAAGCCGGTTCAGCTTC
CAGGTGCTTACAACGTAAACATCAAACTTGATATTACAAGCCACAATGAAGATTATACGATTGTTGAA
CAATATGAACGCGCTGAAGGCCGTCATTCAACTGGCGGAATGGATGAGCTTTACAAATAACGTCTCGG
TTTAAGAGCTATGCTGGAAACAGCATAGCAAGTTTAAATAAGGCTAGTCCGTTATCAACTTGAAAAAG
TGGCACCGAGTCGGTGCTTTTTTTCTGTCTCTTATACACATCTCCGAGCCCACGAGACTAAGGCGAATC
TCGTATGCCGTCTTCTGCTTGCTCGAGGCGTATCTAGACGAGATC

**Table S9. Barcodes used for different samples in MiniSeq**

| Index 1 (i7) | Sequence | Index 2 (i5) | Sequence |
|---|---|---|---|
| N701 | TAAGGCGA | N501 | TAGATCGC |
| N702 | CGTACTAG | N502 | CTCTCTAT |
| N703 | AGGCAGAA | N503 | TATCCTCT |
| N704 | TCCTGAGC | N504 | AGAGTAGA |
| N705 | GGACTCCT | N505 | GTAAGGAG |
| N706 | TAGGCATG | N506 | ACTGCATA |
| N707 | CTCTCTAC | N507 | AAGGAGTA |
| N708 | CAGAGAGG | N508 | CTAAGCCT |
| N709 | GCTACGCT | | |
| N710 | CGAGGCTG | | |
| N711 | AAGAGGCA | | |
| N712 | GTAGAGGA | | |

Selection of barcode combinations for pooling samples follows the "nextera low plex pooling guidelines" of illumina.

**The sequence of the amplicons amplified by one-step PCR in the library for MiniSeq**

AATGATACGGCGACCACCGAGATCTACAC**TAGATCGC(N501)**TCGTCGGCAGCGTCAGATGTGTATAAGAGACAG
CCATTCTACAGTTTATTCTTGACATTGCACTGTCCCCCTGGTATAATAACTATANNNNNNNNNNNNNNNNNNNNGT
TTAAGAGCTATGCTGGAAACAGCATAGCAAGTTTAAATAAGGCTAGTCCGTTATCAACTTGAAAAAGTGGCACCG
AGTCGGTGCTTTTTTTCTGTCTCTTATACACATCTCCGAGCCCACGAGAC**TAAGGCGA(N701)**ATCTCGTATGCCGT
CTTCTGCTTG

Notice: Here we used i5 (N501) and i7 (N701) as example barcodes to show the amplicon sequence. The highlighted 20N represents the base-pairing region of sgRNAs. For full annotation of the amplicon, please refer to "CRISPRi-seq amplicon.dna" file on https://www.veeninglab.com/crispri-seq
